# Spike discharge prediction based on Neuro-fuzzy system

**DOI:** 10.1101/133967

**Authors:** Mahdi Zarei

**Affiliations:** University of California, San Francisco

## Abstract

This paper presents the development and evaluation of different versions of Neuro-Fuzzy model for prediction of spike discharge patterns. We aim to predict the spike discharge variation using first spike latency and frequency-following interval. In order to study the spike discharge dynamics, we analyzed the Cerebral Cortex data of the cat from [29]. Adaptive Neuro-Fuzzy Inference Systems (ANFIS), Wang and Mendel (WM), Dynamic evolving neural-fuzzy inference system (DENFIS), Hybrid neural Fuzzy Inference System (HyFIS), genetic for lateral tuning and rule selection of linguistic fuzzy system (GFS.LT.RS) and subtractive clustering and fuzzy c-means (SBC) algorithms are applied for data. Among these algorithms, ANFIS and GFS.LT.RS models have better performance. On the other hand, ANFIS and GFS.LT.RS algorithms can be used to predict the spike discharge dynamics as a function of first spike latency and frequency with a higher accuracy compared to other algorithms.

## 1 Introduction

Recording action potentials (spikes) from the neural cells makes it possible to investigate their health, stability, and sensitivity[13]. Different characteristics of the electrical activity of neurons can be considered in the study of neural coding. One important concept in this area is spike discharge that is a type of transient waveforms present in the brain activity and includes a high correlation with seizure occurrence [23].

Studies on movement indole illustrated that this process is related to the neuronal discharge [14, 7]. For example, the study on activity of arm-related neurons and their relationship between premotor cortical cell activity and direction of arm movement shows that the cells activity varies in an orderly fashion with the direction of movement [4]. Also, detection of spike discharge in the electroencephalogram is an important way of diagnosis of the disease [23]. Different algorithms like neural networks, logistic regression, and neuro-fuzzy model can be applied for detection of epileptic seizure [23, 19, 25]. There are many similarities between human and animal brain’s neural coding and many studies used animal modeling for investigation the spike discharge ([21, 18, 22, 5]). Johnsen et al. [13] analyzed twenty-six pairs of units recorded from twenty-four retinal ganglion cells in the isolated goldfish retina and examined the cross-correlation histogram for the maintained discharge of each pair of cells. Their results showed that it is unlikely that differences in latency could be attributed to the unequal effectiveness of the stimuli for the two units. Batuev et al. [3] investigated the postsynaptic response of motor cortex neurons of the cat in response to the stimulation of different modalities and showed that it responds with a wide range of peripheral inputs. The electrical changes in the cerebral cortex can correspond with the electric changes in muscle and nerve [1]. The studies of the functional organization of the motor cortex show that this cortical area is composed of modules consisting of columnar aggregates of neurons related to different aspects of the same movement [16]. The current-flow and current-source-density analysis of the direct cortical response in the somatosensory cortex of rats show that the activation and magnitude of direct cortical response depend on stimulus strength and frequency [10].

In this paper, the variation of spike discharge as a function of first spike latency and the frequency-following interval is analyzed. First spike latency is the time delay between stimulus onset and first action potential [8]. Neuro-fuzzy model is a combination of artificial neural network (ANN) and fuzzy logic approaches. It is a powerful tool for dealing with uncertainty and widely used for analyzing electrical activity of neurons. It is widely used for analyzing the electrical activity of the neurons ([9, 11, 20, 24]). The ANFIS method was successfully applied for EEG signals with a high accuracy of the results obtained [9]. A feature extraction method through the time-series prediction based on ANFIS model for brain-computer interface applications has been proposed by Hsu [11]. In this model, ANFISs is used for prediction of time-series for the left and right motor imagery classification, respectively. It is shown that neuro-fuzzy is an accurate model diagnosing epilepsy [24].

Different versions of the neuro-fuzzy model have been used to find the model with higher accuracy. In all models, the spike discharge is considered as an output of the model, while first spike latency and spike frequency are considered as inputs. Using neuro-fuzzy model as a predictor of spike discharge, we are able to use insufficient crisp inputs to make an accurate decision about spike discharge. We used first spike latency and frequency-following interval in the input layer of the neuro-fuzzy system and output was the spike discharge. The structure of this paper is as follows.

First, we discuss spike discharge, latency, and frequency. Section 2 provides a brief description of ANFIS, WM, DENFIS, HyFIS and SBC algorithms and section 3 presents the performance of different neuro-fuzzy algorithms for analysis of cat data.

## 2 Neuro-fuzzy model

Neuro-fuzzy model is a combination of artificial neural networks and fuzzy logic and it uses capabilities of both models. It applies a neural networks structure and at the same time uses *if-then* rules in fuzzy systems. It uses prior knowledge to compute membership function and different learning algorithms of neural networks, including the back-propagation algorithm [27].

The different types of neuro-fuzzy systems used in this paper are as follow:

- Adaptive Neuro-Fuzzy Inference Systems (ANFIS)
- Wang and Mendel (WM)
- Dynamic evolving neural-fuzzy inference system (DENFIS)
- Hybrid neural Fuzzy Inference System (HyFIS) [17]
- genetic for lateral tuning and rule selection of linguistic fuzzy system (GFS.LT.RS) [2]
- subtractive clustering and fuzzy c-means (SBC) [32, 6]

Here we provide a short description each of them.

Adaptive Neuro-Fuzzy Inference Systems (ANFIS) model is a well-known neuro-fuzzy system that implements a Sugeno fuzzy system and uses a t-norm and differentiable membership function [12, 26]. For a system with two rules, we can build the following neuro-fuzzy structure.

For given two inputs *x*_0_ and *y*_0_ and corresponding linguistic labels *A*_*i*_ and *B*_*i*_, each neuron in the first layer of neuro-fuzzy model transmit the crisp signal to the next layer (Algorithm 1).

### Algorithm 1 ANFIS model

This algorithm has two main stages, the forward and backward steps. The forward step has five layers as follows:

- First layer maps the crisp inputs using bell-shaped membership function as follows:

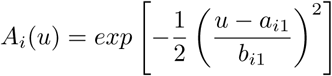

and

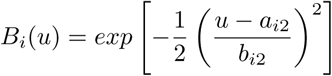

where {a_*i*1_, a_*i*2_, b_*i*1_, b_*i*2_} is the parameters set.
- The second layer is responsible for fuzzification and each neuron in this layer determines the fuzzy degree received crisp input.

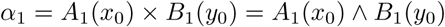

and

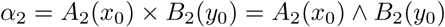
- Neurons in the third layer correspond to fuzzy rules and receive inputs from fuzzification neurons in the second layer. The outputs of layer 3 are as follow:

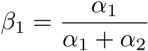

and

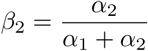
- Layer 4 or output membership layer combine all its inputs by using the fuzzy operation union

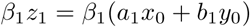

and

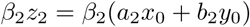
- The last layer is responsible for Defuzzification.

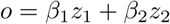

In the backward process, the errors are propagated backward and the parameters are updated by gradient descent technique.

Wang and Mendel (WM) model is another type of neuro-fuzzy system that developed by Wang and Mendel [31] that has high performance for regression tasks. First, it divides input and outputs into the fuzzy region and assigns a membership function to each region. Then finds a rule for each pair of input data. In the next step, a degree is assigned to each rule. After assigning degrees, they are combined. The final rule is obtained after deleting redundant rules. Algorithm 2 provides more details about WM algorithm.

### Algorithm 2 Wang and Mendel (WM)

Division numerical input and output data spaces into fuzzy regions

Generate fuzzy IF-THEN rules covering the training data Determining a degree for each rule

Eliminating redundant rules and obtaining a final rule base

Dynamic evolving neural-fuzzy inference system (DENFIS) is another fuzzy inference system that developed by Kasabov et al. [15]. The output of the system is based on m-most activated fuzzy rules and evolving clustering method is applied to determine the cluster center (Algorithm 3).

### Algorithm 3 Dynamic evolving neural-fuzzy inference system (DENFIS) model

Choose cluster center from training data

Determine the cluster centers using the evolving clustering method partition the input space and to find optimal parameters on the consequent part Update the parameters on the consequent part

Hybrid neural Fuzzy Inference System (HyFIS) has two general steps for learning [17]. In the first step, the Wang and Mendel is used for knowledge acquisition. In the second step, the input vector is propagated forward in the network and parameter updating is performed using backpropagating the error using a gradient descending approach [28].

### Algorithm 4 Hybrid neural Fuzzy Inference System (HyFIS)

Uses the techniques of Wang and Mendel to acquire the knowledge

Use gradient descent-based to learn parameters of the structure

**GFS.LT.RS**: GFS.LT.RS is proposed by R. Alcala et al. [2] that performs an evolutionary lateral tuning of membership functions in constructing FRBS model to obtain higher accurate linguistic models (algorithm 5).

### Algorithm 5 genetic for lateral tuning and rule selection of linguistic fuzzy system (GFS.LT.RS)

Uses the Wang and Mendel to to construct the population

Evaluate the chromosome using Mean square error

Minimize the number of rules

Subtractive clustering and fuzzy c-means (SBC) [32, 6] is checking each data point’s distance from all other data points to find the cluster centers. More details about SBC algorithm is provided in the Algorithm 6

### Algorithm 6 Subtractive clustering and fuzzy c-means (SBC)

Use subtractive clustering method to obtain the cluster centeres (generating the rules)

Choose the highest potential as the cluster centere

Update the potential of each data point

Optimise the cluster centers using fuzzy c-means

## 3 computational results

To verify the effectiveness of the neuro-fuzzy algorithms we carried out a number of numerical experiments with the cortex of the somatosensory/motor system of the Cat data set on a PC with Processor Intel(R) Core(TM) i5-3470S CPU 2.90 GHz and 8 GB RAM running under Windows XP. The cortex of the somatosensory/motor system of the Cat data is publicly available from [29]. This data is based on recording neurons of extracellularly in post cruciate cerebral cortex of cats. It is neuronal responsiveness of each of the four paws to strong cortical surface stimulation to understand facilitatory and inhibitory modulation of wide-field neurons by small-field neurons. Two groups of data from the Cerebral Cortex of the Cat data sets are considered for evaluation of the algorithms: Contralateral Forepaw (CF) Cortex (Chloralose) and Contralateral Hindpaw (CH) Cortex (Chloralose). The Contralateral Forepaw (CF) Cortex (Chloralose) is based on the measurements of 4,272 neurons, but Contralateral Hindpaw (CH) Cortex (Chloralose) contains data of 991 neurons. Various versions of neuro-fuzzy algorithms from R package are used to evaluate the algorithms’ error for each data. The R Project for Statistical Computing is an environment for statistical computing and graphics that contains comprehensive libraries of machine learning and statistical analysis applications that are available on [30].

### 3.1 Results of Forepaw (CF) Cortex (Chloralose) analysis

The Ipsilateral and Contralateral data from Forepaw Cortex data are considered for analysis. The results of application of neuro-fuzzy algorithm to the Ipsilateral Forepaw Cortex data are presented on the Figures 3.1-3.1. The first spike latency and ipsilateral forepaw frequency following interval (msec) are used as inputs, while mean spikes per discharge is used as output of the model. Each figure contains the actual spikes per discharge value that is computed using neuro-fuzzy algorithm. Also, some statistics about the analysis is illustrated in each figure. The results show that the smallest Root Mean Square Error (RMSE) is obtained by HYFIS algorithm (RMSE=1.34) and the biggest RMSE is obtained by WM model (RMSE=2.72).

**Figure 1:**
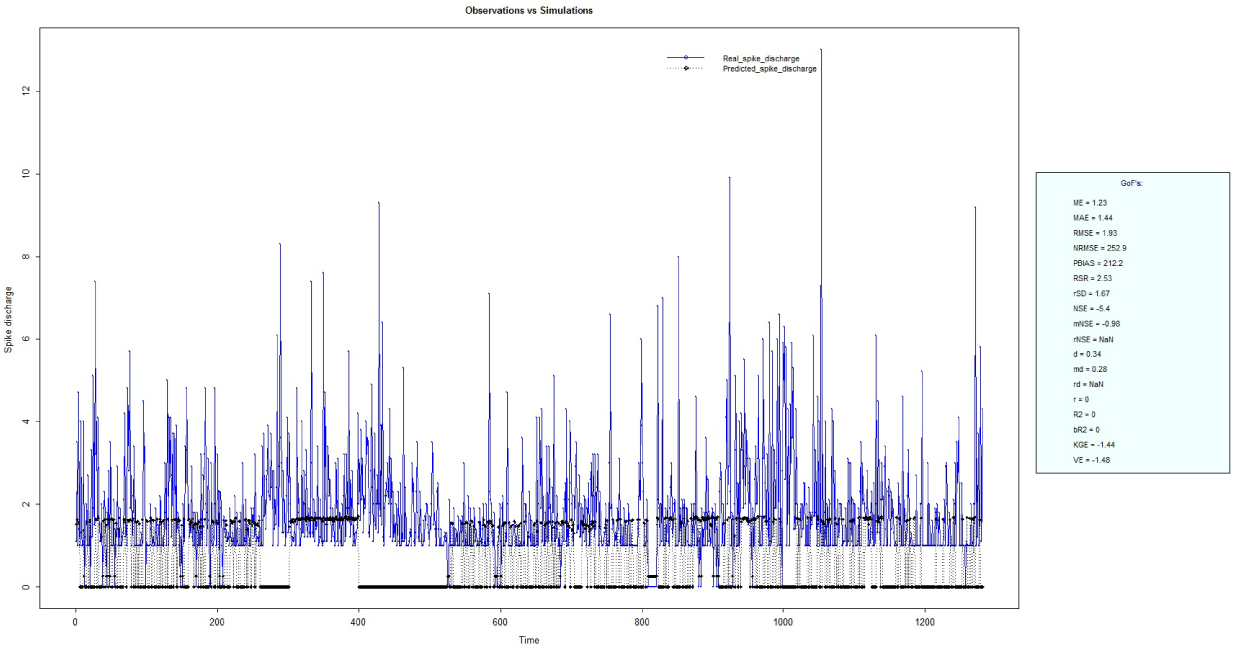
Spike Discharge prediction for cat Ipsilateral Forepaw Cortex using ANFIS algorithm.

**Figure 2:**
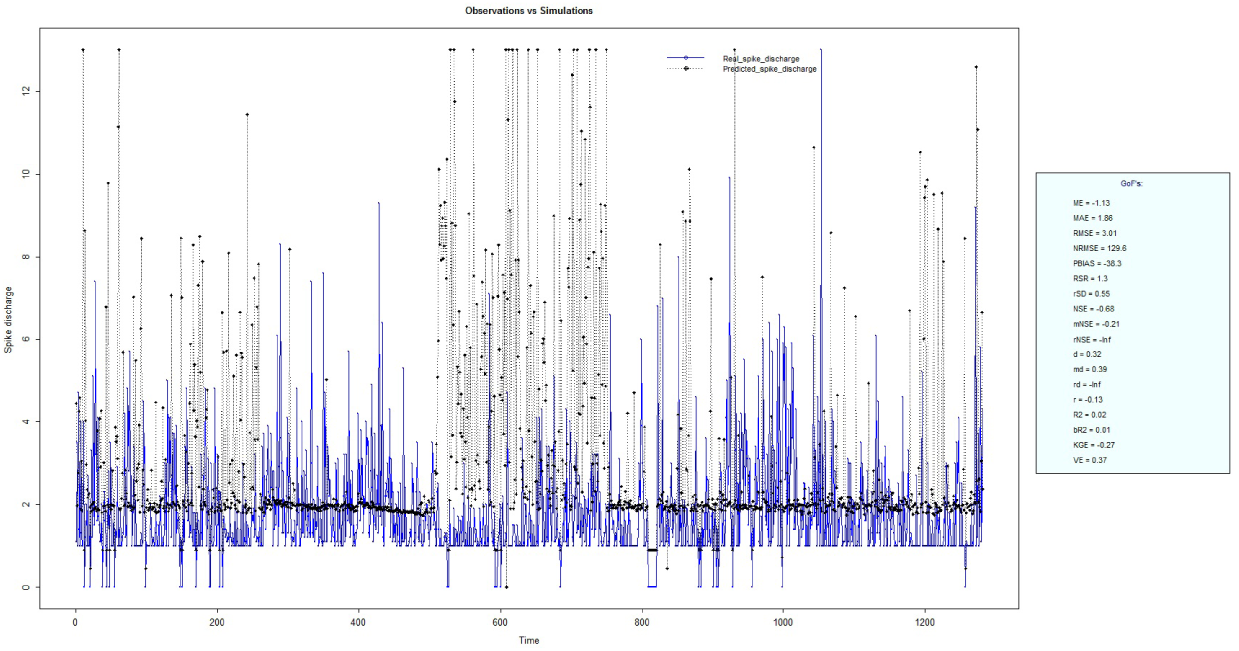
Spike Discharge prediction for cat Ipsilateral Forepaw Cortex using Denfis algorithm. Figures 3.1 -3.1 present the results of application for neuro-fuzzy algorithm for the Contralateral Forepaw Cortex data. Again the 1st spike latency and ipsilateral forepaw frequency following interval (msec) are used as inputs and mean spikes per discharge is used as output of the model. The results demonstrate that the smallest Root Mean Square Error (RMSE) is obtained using HYFIS algorithm (RMSE=0.93) and the biggest RMSE is obtained by WM model (RMSE=4.27).

### 3.2 Results of Hindpaw Cortex (Chloralose) data analysis

The Hindpaw Cortex is divided into two parts: the Contralateral Forepaw Cortex and Ipsilateral Hindpaw Cortex. Then different neuro-fuzzy algorithms have been applied to them. Figures 3.1-3.1 present the results of application of neuro-fuzzy algorithm for the Contralateral Forepaw Cortex data. The best RMSE is obtained using GFS LT RS (RMSE=2.06), the smallest Root Mean Square Error (RMSE) is obtained using HYFIS algorithm (RMSE=0.93), and the biggest RMSE is btained by WM model (RMSE=4.27).

**Figure 3:**
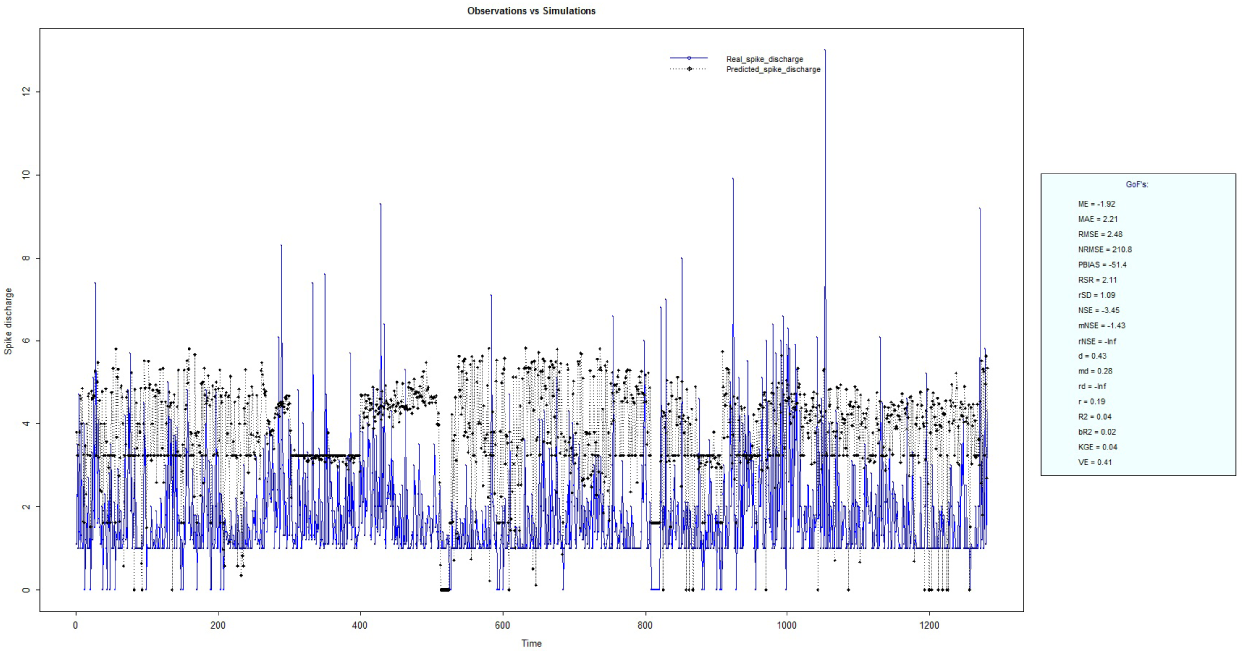
Spike Discharge prediction for cat Ipsilateral Forepaw Cortex using GFS LT RS algorithm.

**Figure 4:**
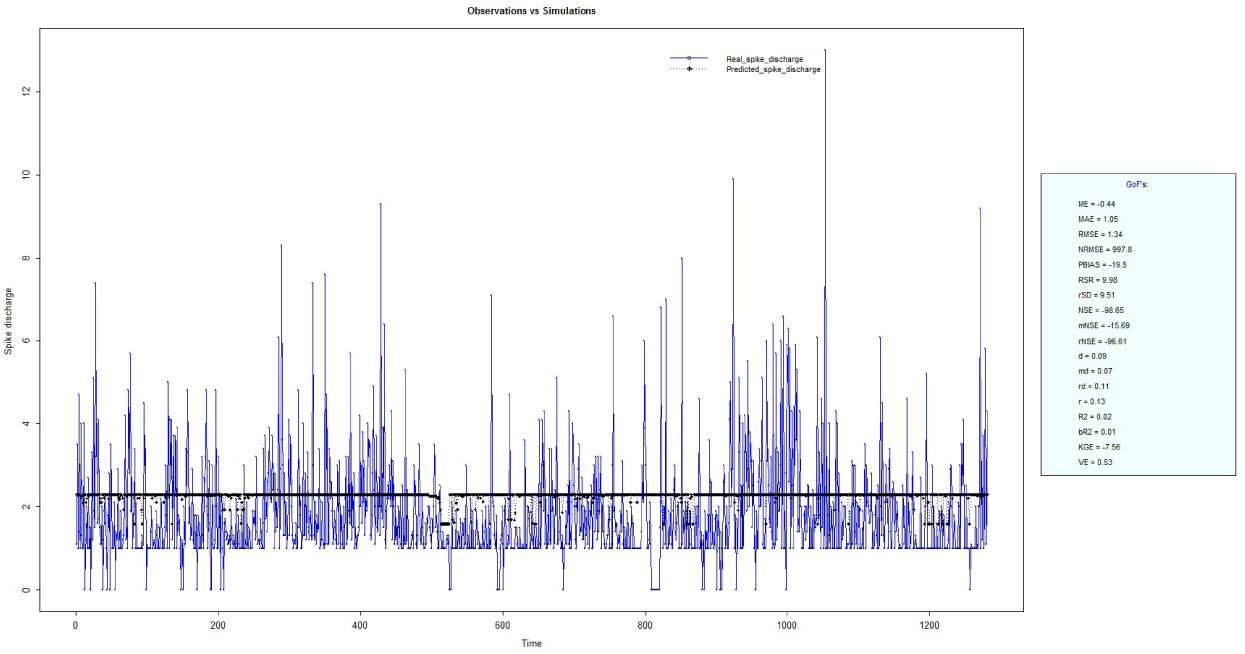
Spike Discharge prediction for cat Ipsilateral Forepaw Cortex using HYFIS algorithm.

**Figure 5:**
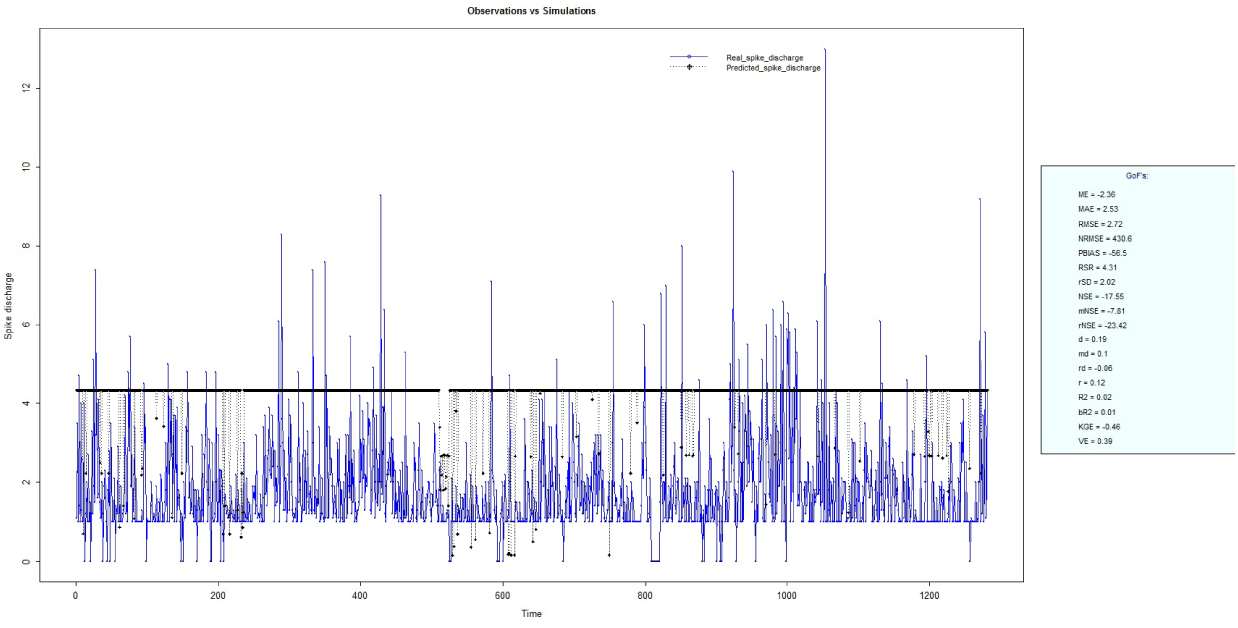
Spike Discharge prediction for cat Ipsilateral Forepaw Cortex using WM algorithm. Results of application of the algorithms to the Ipsilateral Hindpaw Cortex data are presented in figures 3.2-3.2. The WM algorithm provides better accuracy compared with other algorithm (RMSE=2.73)

**Figure 6:**
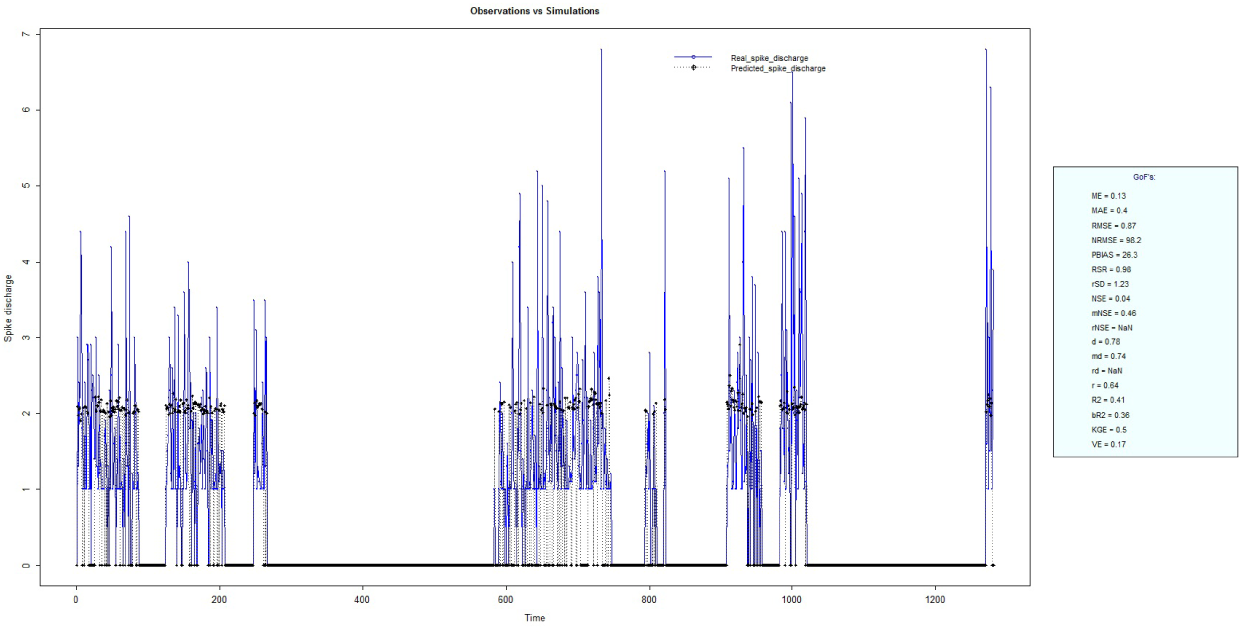
Spike Discharge prediction for cat Contralateral Forepaw Cortex using ANFIS algorithm.

**Figure 7:**
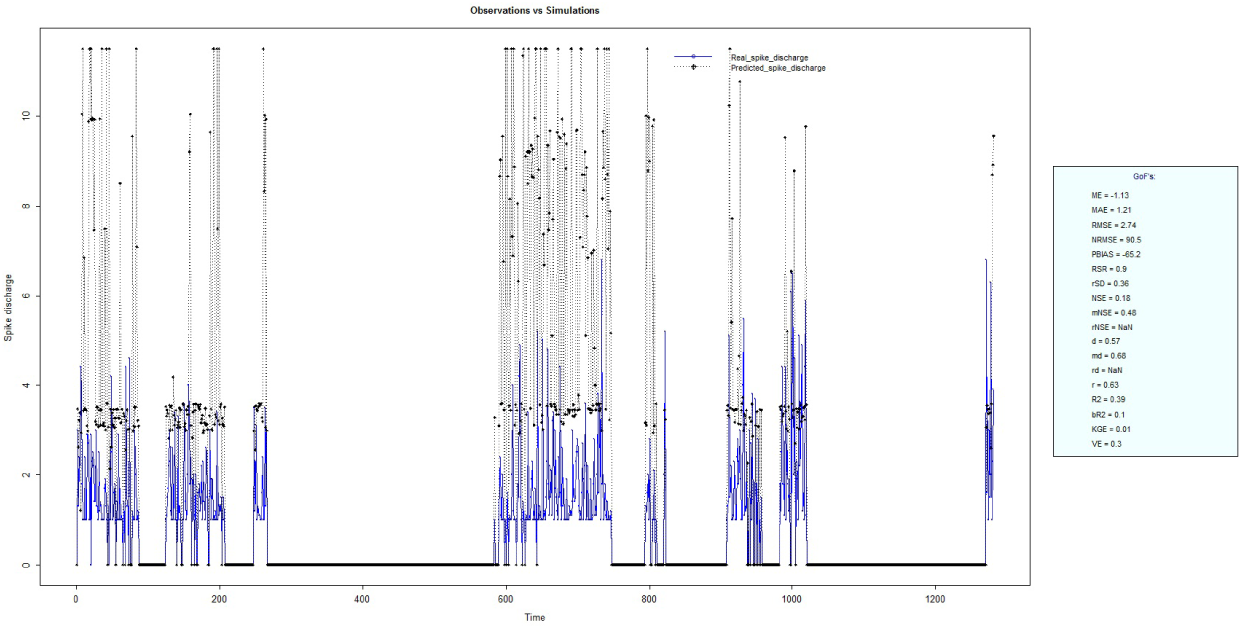
Spike Discharge prediction for cat Contralateral Forepaw Cortex using Denfis algorithm.

**Figure 8:**
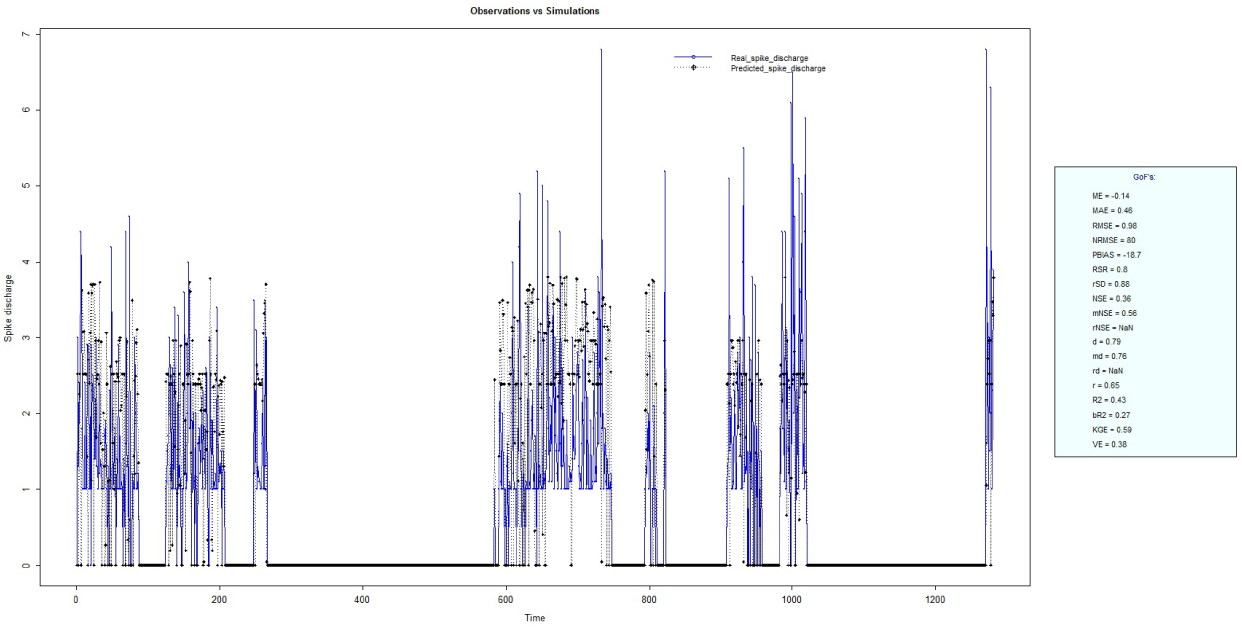
Spike Discharge prediction for cat Contralateral Forepaw Cortex using GFS LT RS algorithm.

**Figure 9:**
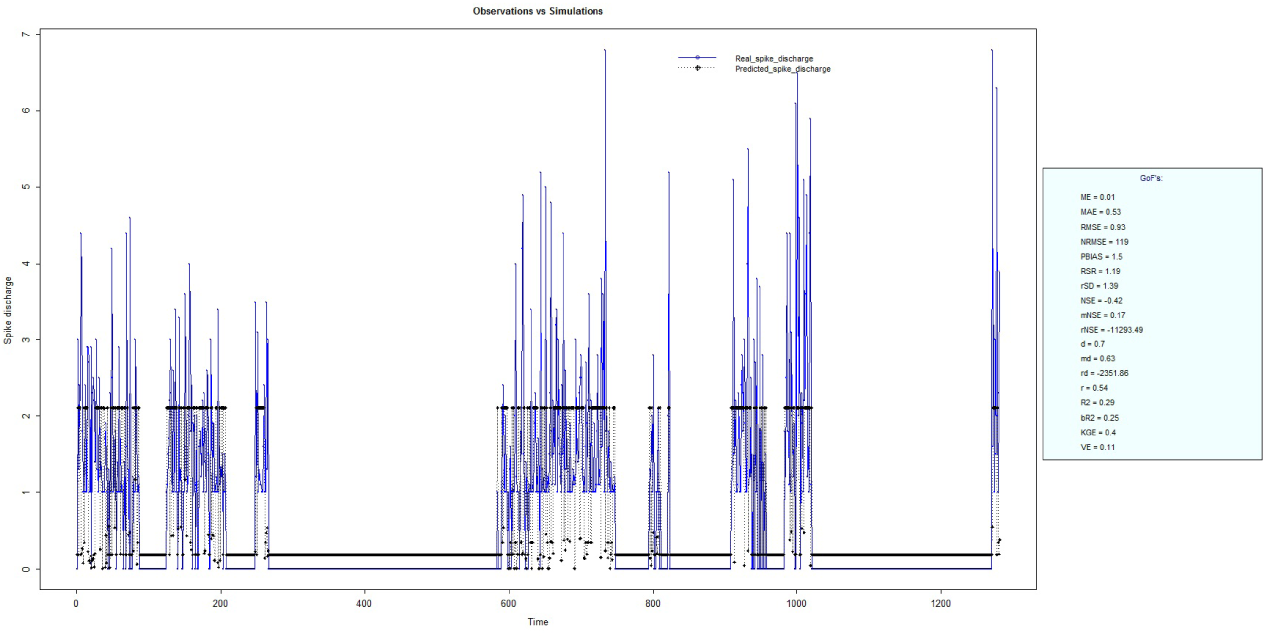
Spike Discharge prediction for cat Contralateral Forepaw Cortex using HYFIS algorithm.

**Figure 10:**
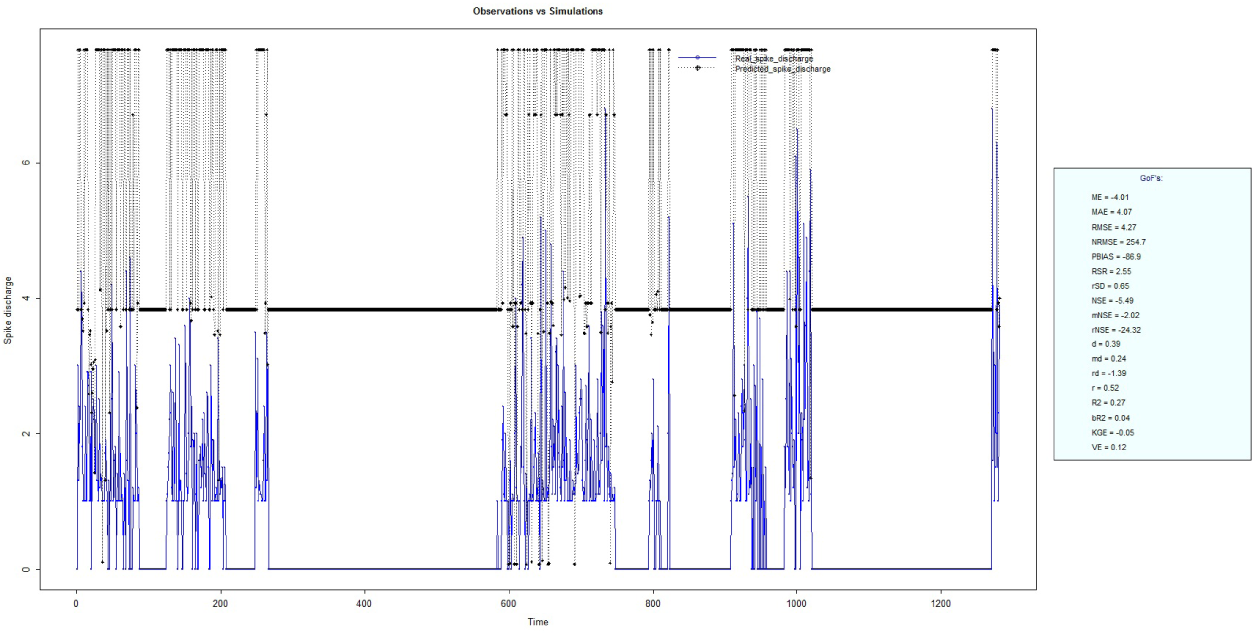
Spike Discharge prediction for cat Contralateral Forepaw Cortex using WM algorithm.

**Figure 11:**
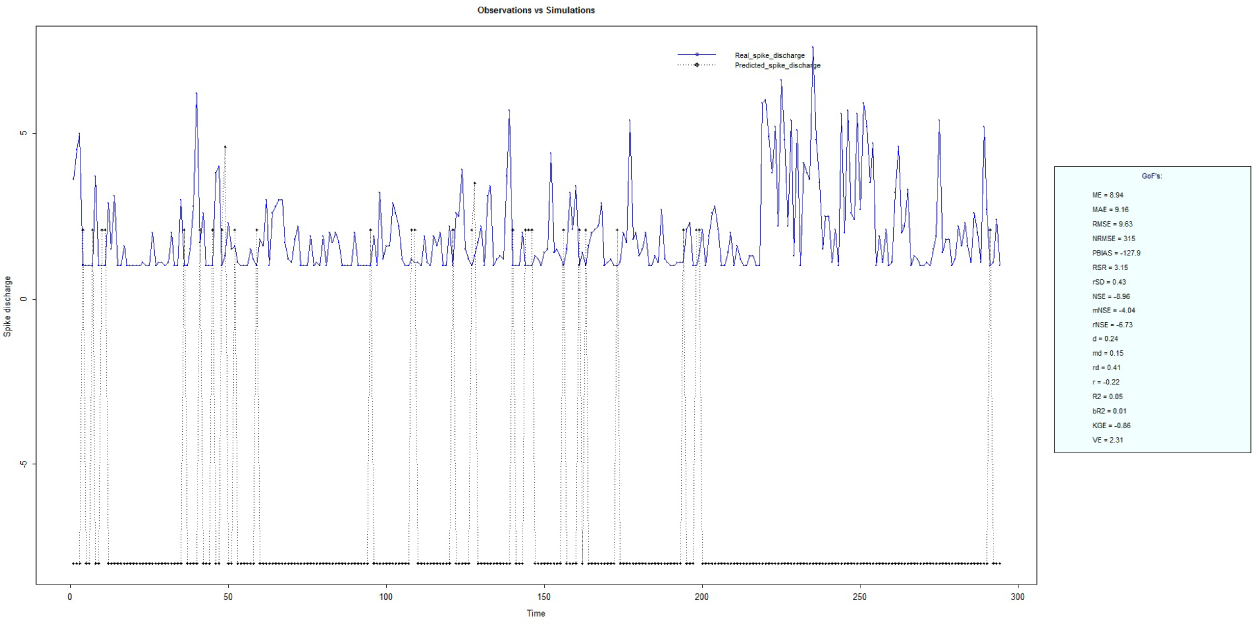
Spike Discharge prediction for cat Contralateral Hindpaw Cortex using ANFIS algorithm.

**Figure 12:**
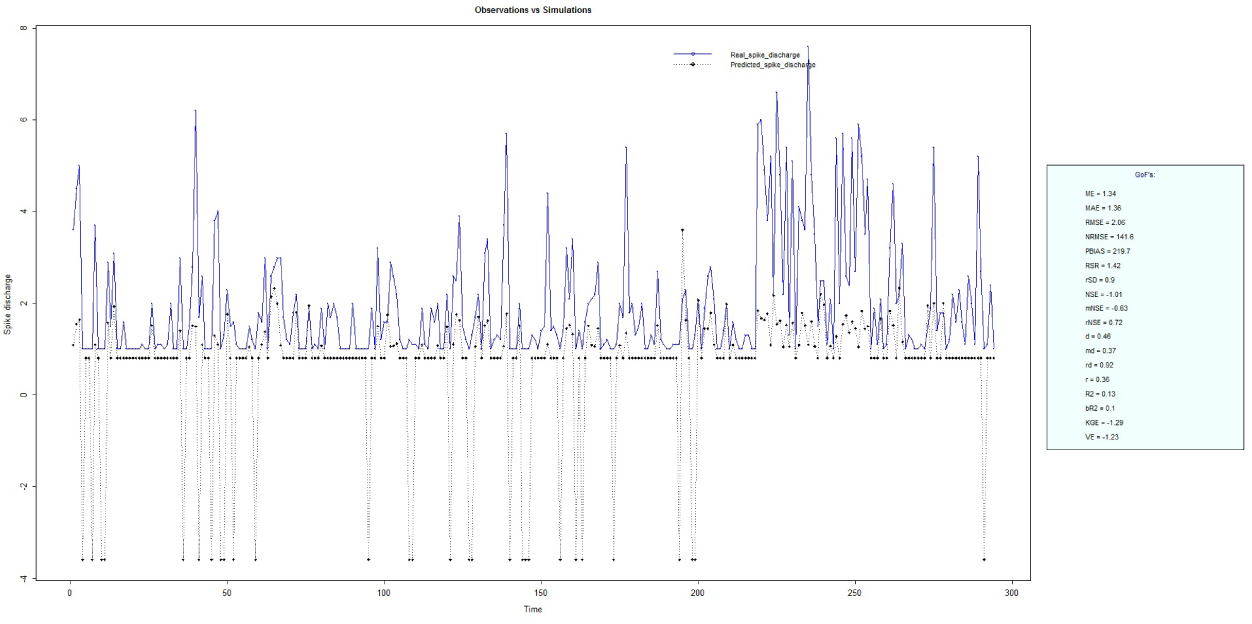
Spike Discharge prediction for cat Contralateral Hindpaw Cortex using Denfis algorithm.

**Figure 13:**
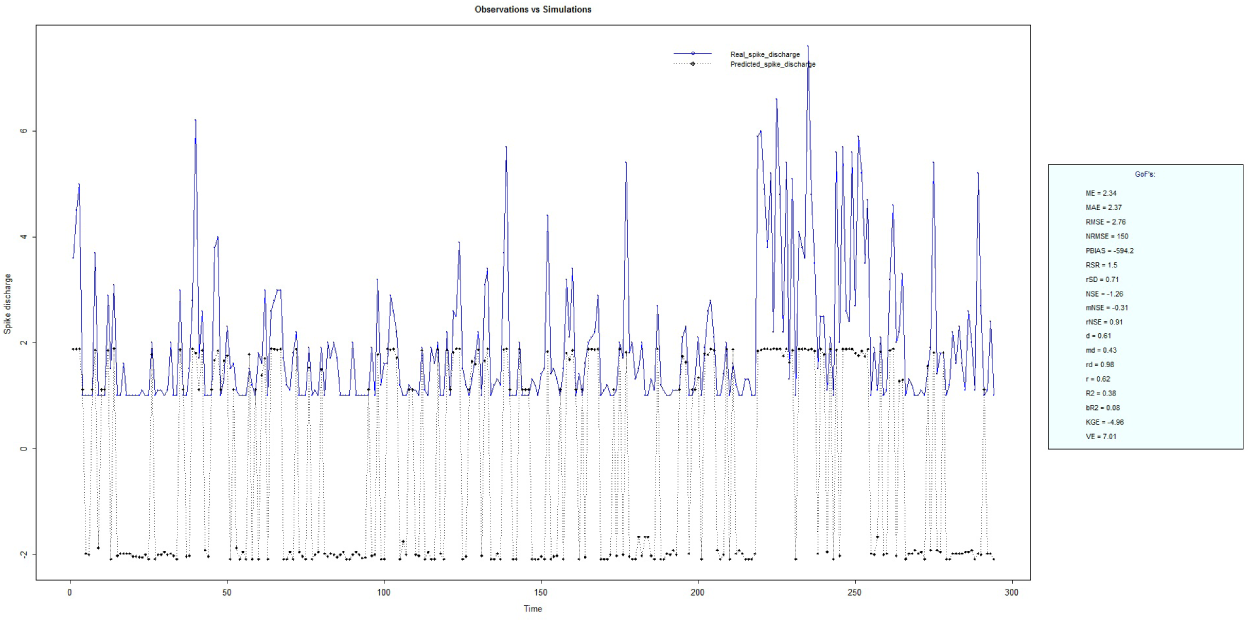
Spike Discharge prediction for cat Contralateral Hindpaw Cortex using SBC algorithm.

**Figure 14:**
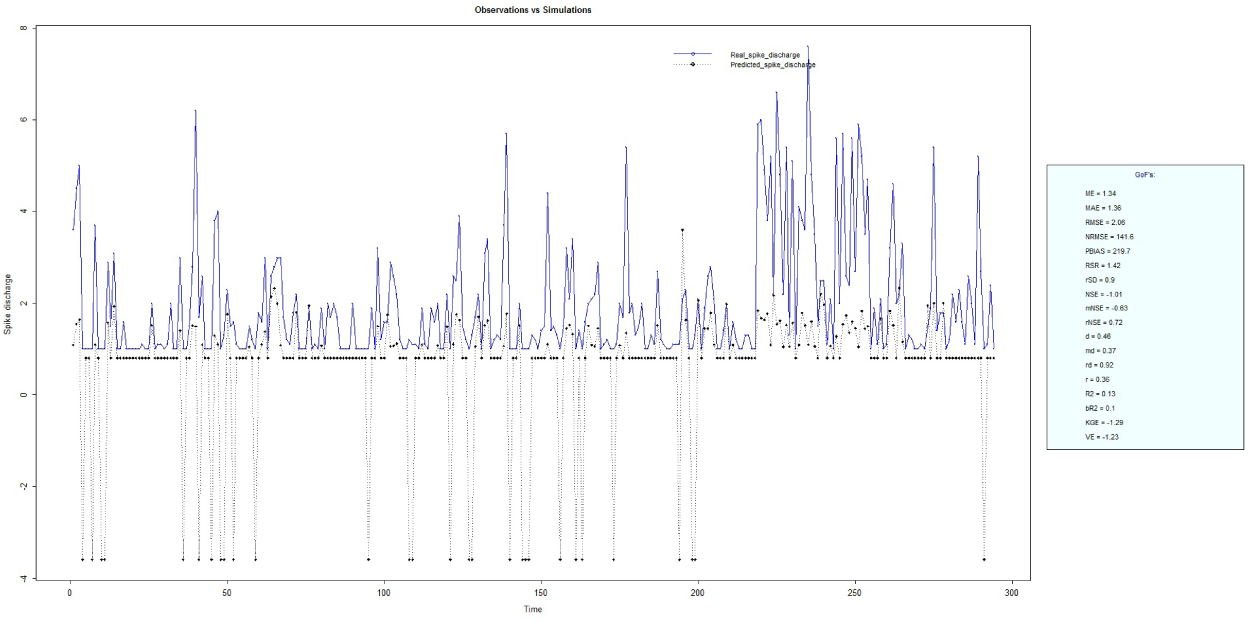
Spike Discharge prediction for cat Contralateral Hindpaw Cortex using GFS LT RS algorithm.

**Figure 15:**
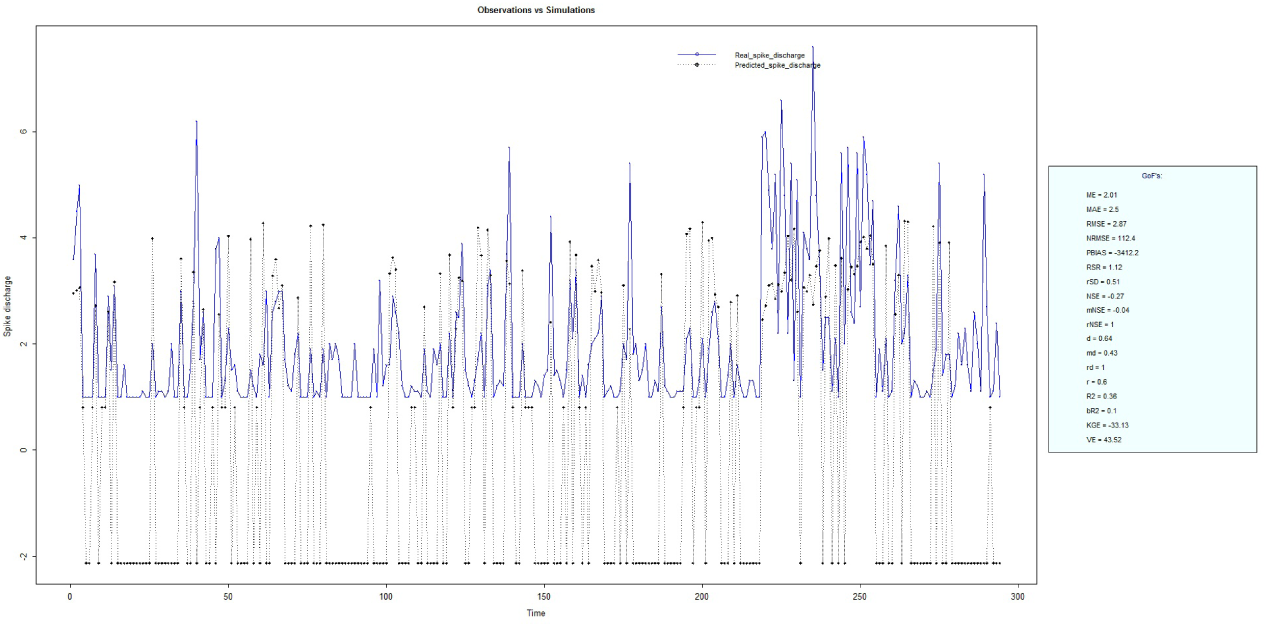
Spike Discharge prediction for cat Contralateral Hindpaw Cortex using WM algorithm.

**Figure 16:**
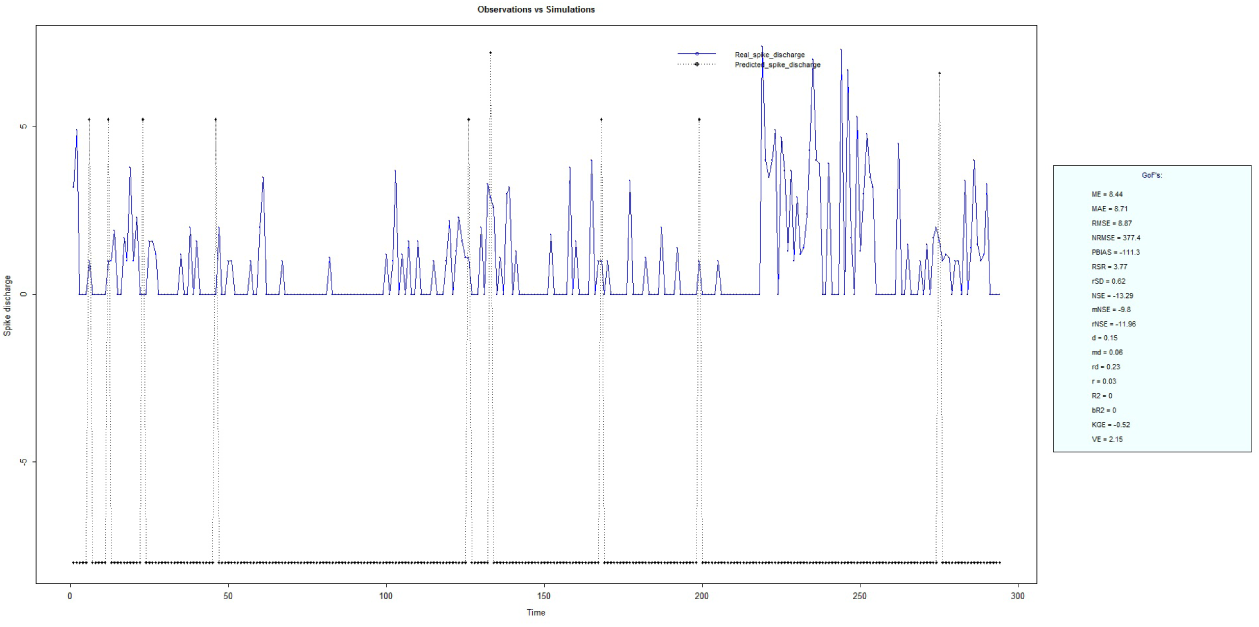
Spike Discharge prediction for cat Ipsilateral Hindpaw Cortex using ANFIS algorithm.

**Figure 17:**
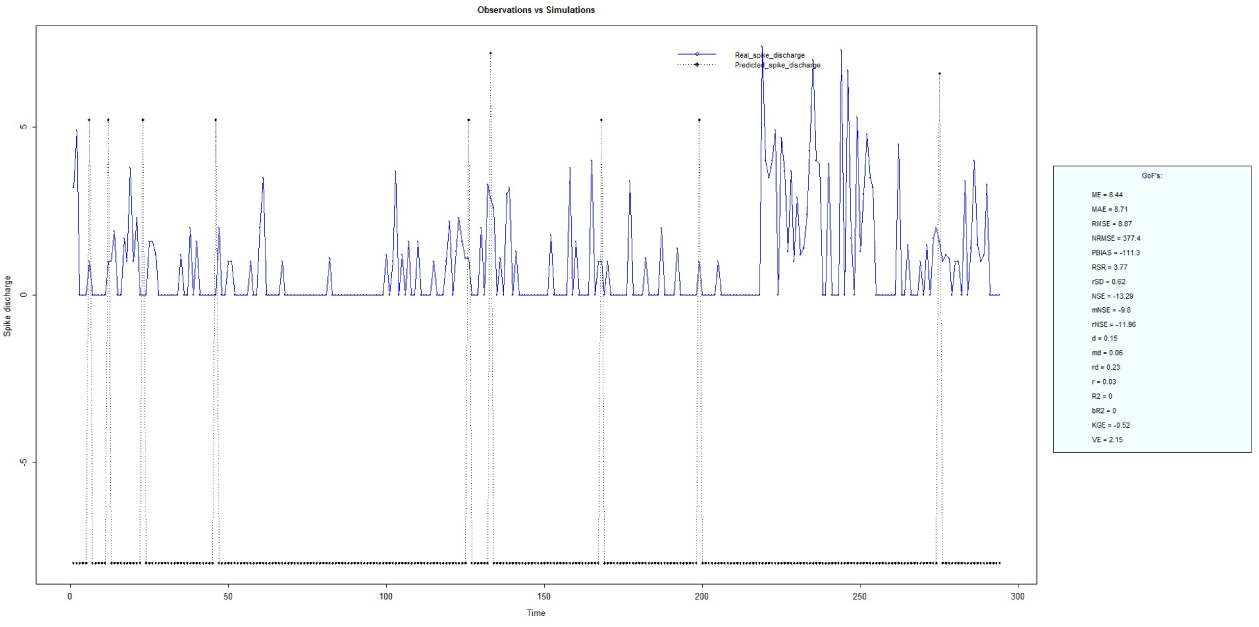
Spike Discharge prediction for cat Ipsilateral Hindpaw Cortex using Denfis algorithm.

**Figure 18:**
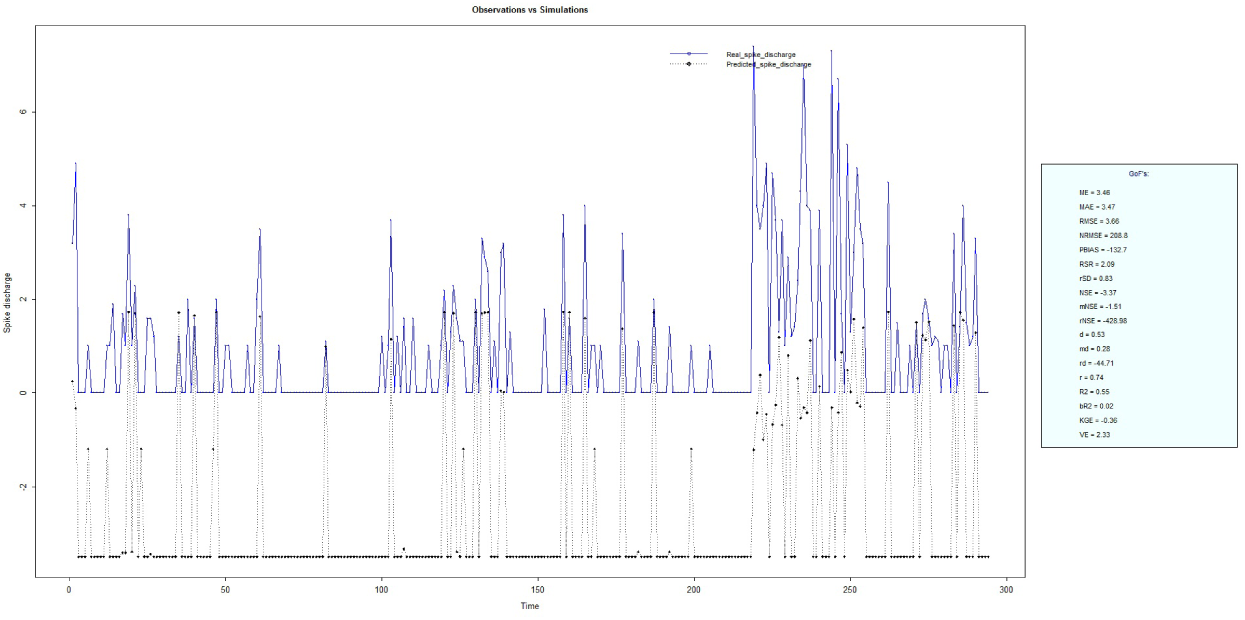
Spike Discharge prediction for cat Ipsilateral Hindpaw Cortex using SBC algorithm.

**Figure 19:**
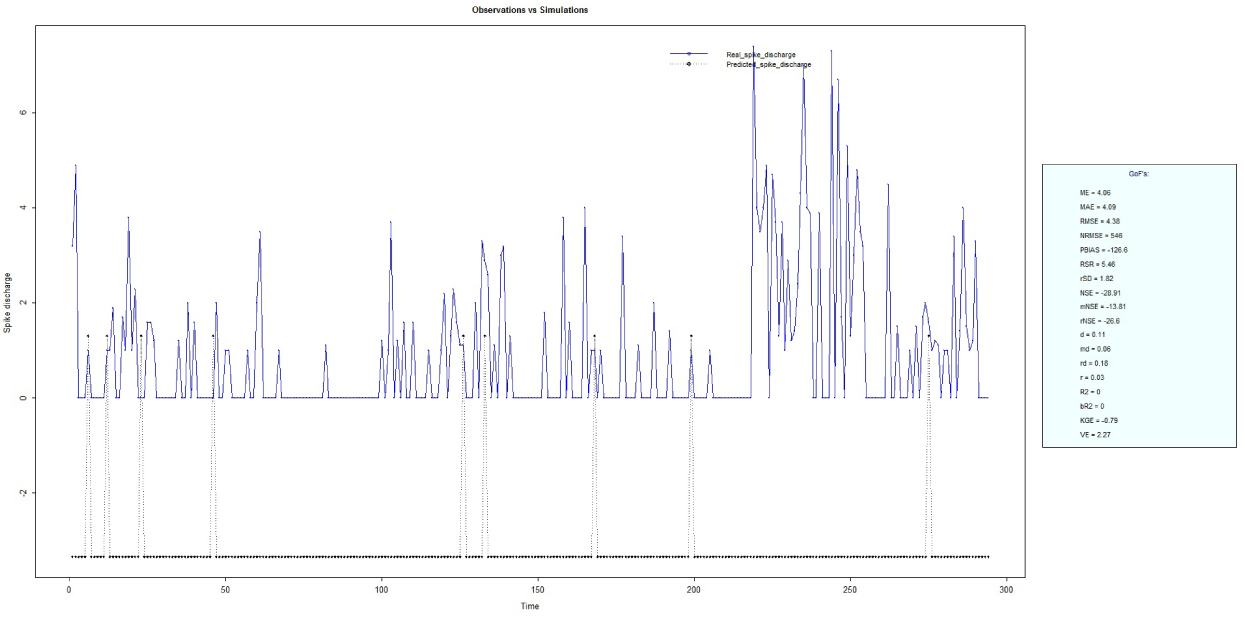
Spike Discharge prediction for cat Ipsilateral Hindpaw Cortex using GFS LT RS algorithm.

**Figure 20:**
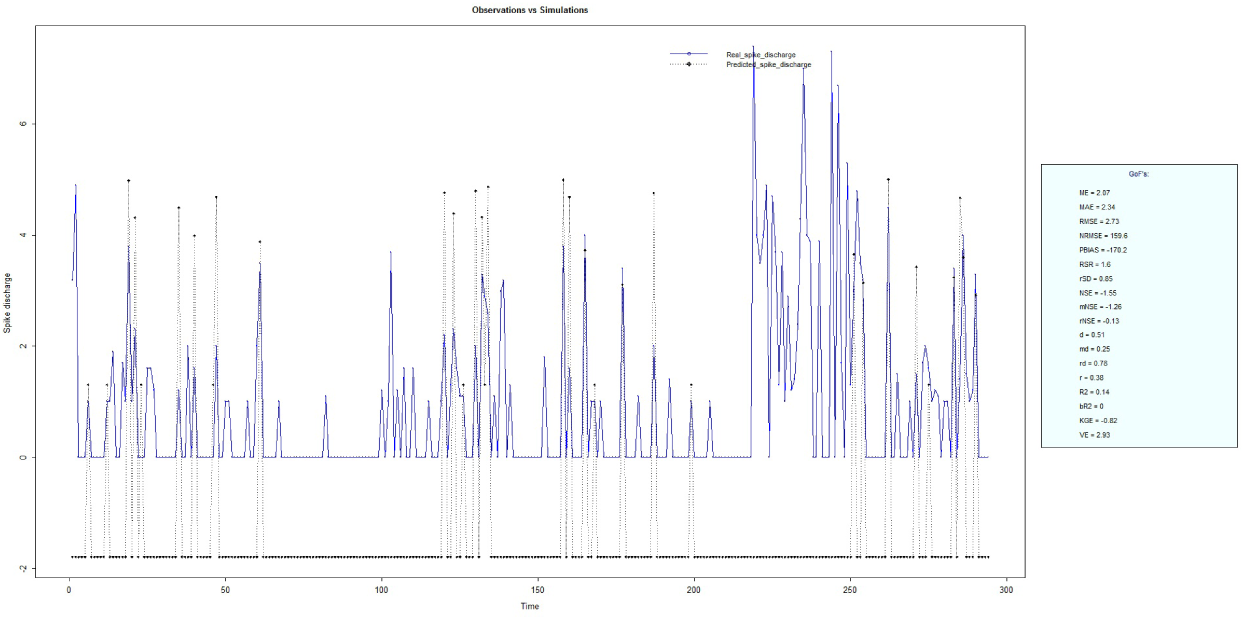
Spike Discharge prediction for cat Ipsilateral Hindpaw Cortex using WM algorithm.

## Conclusion

In this section we presented the development and evaluation of different versions of adaptive neurofuzzy model including Adaptive Neuro-Fuzzy Inference Systems, Wang and Mendel, Dynamic evolving neural-fuzzy inference system, Hybrid neural Fuzzy Inference System, genetic for lateral tuning and rule selection of linguistic fuzzy system and subtractive clustering and fuzzy c-means algorithms for prediction of Spike discharge. Results reveal that Spike discharge can be predicted using the neuro-fuzzy model where first spike latency and frequency-following interval are the inputs and spike discharge is the output of the model.

## References

[1] Adrian, E. D. The spread of activity in the cerebral cortex. The Journal of Physiology 88, 2 (1936), 127–161.

[2] Alcala, R., Alcala-Fdez, J., and Herrera, F. A proposal for the genetic lateral tuning of linguistic fuzzy systems and its interaction with rule selection. Fuzzy Systems, IEEE Transactions on 15, 4 (2007), 616–635.

[3] Batuev, A., and Pirogov, A. Postsynaptic responses of motor cortex neurons of cats to sensory stimijlaation of different modalities. Acta Neurobiologiae Experimentalis 34 (1974), 317–321.

[4] Caminiti, R., Johnson, P., Galli, C., Ferraina, S., Burnod, Y., and Urbano, A. Making arm movements within different parts of space: the premotor and motor cortical representation of a coordinate system for reaching to visual targets. The Journal of Neuroscience 11, 5 (1991), 1182–1197.

[5] Cavelier, P., and Bossu, J.-L. Dendritic low-threshold ca2+ channels in rat cerebellar purkinje cells: possible physiological implications. The Cerebellum 2, 3 (2003), 196–205.

[6] Chiu, S. Method and software for extracting fuzzy classification rules by subtractive clustering. In Fuzzy Information Processing Society, 1996. NAFIPS., 1996 Biennial Conference of the North American (1996), IEEE, pp. 461–465.

[7] Fu, Q.-G., Flament, D., Coltz, J., and Ebner, T. Relationship of cerebellar purkinje cell simple spike discharge to movement kinematics in the monkey. Journal of Neurophysiology 78, 1 (1997), 478–491.

[8] Gollisch, T., and Meister, M. Rapid neural coding in the retina with relative spike latencies. Science 319, 5866 (2008), 1108–1111.

[9] Güler, I., and Übeyli, E. D. Adaptive neuro-fuzzy inference system for classification of eeg signals using wavelet coefficients. Journal of Neuroscience Methods 148, 2 (2005), 113–121.

[10] Harding, G. The currents that flow in the somatosensory cortex during the direct cortical response. Experimental Brain Research 90, 1 (1992), 29–39.

[11] Hsu, W.-Y. Eeg-based motor imagery classification using neuro-fuzzy prediction and wavelet fractal features. Journal of Neuroscience Methods 189, 2 (2010), 295–302.

[12] Jang, J. S. R. Anfis: adaptive-network-based fuzzy inference system. Systems, Man and Cybernetics, IEEE Transactions on 23, 3 (1993), 665–685.

[13] Johnsen, J. A., and Levine, M. W. Correlation of activity in neighbouring goldfish ganglion cells: relationship between latency and lag. The Journal of Physiology 345, 1 (1983), 439–449.

[14] Kalaska, J. What parameters of reaching are encoded by discharges of cortical cells. Motor control: Concepts and Issues(1991), 307–330.

[15] Kasabov, N. K., and Song, Q. Denfis: dynamic evolving neural-fuzzy inference system and its application for time-series prediction. Fuzzy Systems, IEEE Transactions on 10, 2 (2002), 144–154.

[16] Keller, A. Intrinsic synaptic organization of the motor cortex. Cerebral Cortex 3, 5 (1993), 430–441.

[17] Kim, J., and Kasabov, N. Hyfis: adaptive neuro-fuzzy inference systems and their application to nonlinear dynamical systems. Neural Networks 12, 9 (1999), 1301–1319.

[18] Mariño, J., Canedo, A., and Aguilar, J. Sensorimotor cortical influences on cuneate nucleus rhythmic activity in the anesthetized cat. Neuroscience 95, 3 (1999), 657–673.

[19] Pang, C. C., Upton, A. R., Shine, G., and Kamath, M. V. A comparison of algorithms for detection of spikes in the electroencephalogram. Biomedical Engineering, IEEE Transactions on 50, 4 (2003), 521–526.

[20] Pohl, V., and Fahr, E. Neuro-fuzzy recognition of k-complexes in sleep eeg signals. In Engineering in Medicine and Biology Society, 1995., IEEE 17th Annual Conference (1995), vol. 1, IEEE, pp. 789–790.

[21] Prince, D., and Farrell, D. Centrencephalic spike-wave discharges following parenteral penicillin injection in cat. In Neurology (1969), vol. 19, LIPPINCOTT WILLIAMS & WILKINS 227 EAST WASHINGTON SQ, PHILADELPHIA, PA 19106, p. 309.

[22] Sitnikova, E. Thalamo-cortical mechanisms of sleep spindles and spike–wave discharges in rat model of absence epilepsy (a review). Epilepsy Research 89, 1 (2010), 17–26.

[23] Subasi, A. Automatic detection of epileptic seizure using dynamic fuzzy neural networks. Expert Systems with Applications 31, 2 (2006), 320–328.

[24] Subasi, A. Application of adaptive neuro-fuzzy inference system for epileptic seizure detection using wavelet feature extraction. Computers in Biology and Medicine 37, 2 (2007), 227–244.

[25] Subasi, A., and Erçelebi, E. Classification of eeg signals using neural network and logistic regression. Computer Methods and Programs in Biomedicine 78, 2 (2005), 87–99.

[26] Takagi, T., and Sugeno, M. Fuzzy identification of systems and its applications to modeling and control. Systems, Man and Cybernetics, 1 (1985), 116–132.

[27] Tsoukalas, L. H., and Uhrig, R. E. Fuzzy and neural approaches in engineering. John Wiley & Sons, Inc., 1996.

[28] Tung, S. W., Quek, C., and Guan, C. T2-hyfis-yager: Type 2 hybrid neural fuzzy inference system realizing yager inference. In Fuzzy Systems, 2009. FUZZ-IEEE 2009. IEEE International Conference on (2009), IEEE, pp. 80–85.

[29] http://oto2.wustl.edu/bbears/arnie/catcrtx.htmu/bbears/arnie/catcrtx.htm.

[30] http://www.r-project.org/.

[31] Wang, L.-X., and Mendel, J. M. Generating fuzzy rules by learning from examples. Systems, Man and Cybernetics, IEEE Transactions on 22, 6 (1992), 1414–1427.

[32] Yager, R. R., and Filev, D. P. Generation of fuzzy rules by mountain clustering. Journal of Intelligent and Fuzzy Systems 2, 3 (1994), 209–219.

